# A motif for domain-specific analysis applets that are easy to learn, reuse, test, and to compose into pipelines: application to vision science

**DOI:** 10.64898/2026.04.27.721136

**Authors:** Avraham A. Lepsky, Madeline K. Severson, Ruyang Wang, Xinyu Cheng, Ricardo L. Rodriguez, Rui Gong, Stephen D. Van Hooser

**Affiliations:** Brandeis University, Department of Biology, Waltham, MA, USA

## Abstract

Scientific progress depends on the analysis of primary data, yet the small, domain-specific programs that perform most scientific analyses are typically poorly documented, narrowly tested, and difficult to reuse outside the lab that created them. General-purpose pipeline tools address the problem of running steps in order but do not enforce documentation, testing, or standardized outputs. We describe a motif for building domain-specific analysis applets, which we call *calculators*, that constrains developer choices in order to produce code that is readable, tested, and reusable almost as a byproduct of following the template. *Calculators* operate on a typed, searchable database of documents, eliminating the need to explicitly wire inputs and outputs together; instead, each *calculator* searches the database for documents it can operate on and adds its results as new typed documents. *Calculators* must provide documentation in a standard location, self-tests that can be run and inspected interactively, adjustable input parameters, a single well-defined output document type, and a default plotting method. Sets of *calculators* compose naturally into pipelines whose outputs satisfy FAIR principles at every stage. We demonstrate the motif by implementing *calculators* for common analyses in vision science, including orientation and direction selectivity, contrast tuning, spatial and temporal frequency tuning, speed tuning, and Hartley reverse correlation. These *calculators* have been used in published work and are in active use across collaborating laboratories. We discuss the design principles of the motif, its advantages and limitations, and its applicability to domain-specific computation across neuroscience and beyond.

**Significance Statement:** Scientists often must write small programs to analyze their own data. These programs are usually poorly documented, lightly tested, and hard for other labs to reuse. Mistakes in this kind of code have even caused well-known papers to be retracted. We describe a simple pattern for writing these programs, which we call a calculator. The pattern requires the programmer to include clear documentation, built-in tests, adjustable settings, and a standard form of output. Calculators work by searching a shared database for data they know how to handle, so many calculators can be chained together into a pipeline without extra setup. We show how this works by building calculators for common visual neuroscience analyses that other labs are already using.

## 1 INTRODUCTION

Scientific advances depend critically on the analysis of primary data. In vision science and in neuroscience in general, the primary data are complex. Correlating visual stimuli with the responses of the nervous system, such as intracellular voltage records, calcium signals, extracellular spike recordings, or fMRI BOLD signals has required substantial software development on the part of individual researchers and labs.

The normal development of robust software requires several processes (AppJungle.NET LLC [2023]), including deciding what the software will do (scoping), user experience design, software development, documentation, testing and quality assurance, and customer support. Even small software companies have separate teams for these activities. Software that serves very large user bases can generate revenue that supports these activities. By contrast, scientists who create software to analyze specialized data, such as physiological recordings of the visual system, have precious little time to dedicate to these activities. Despite good intentions, investigators develop custom code that is, in general, poorly tested and has very limited applicability, often only operating on a particular data format and dataset organization (Herndon [2014], McElreath [2023], Huang and Lapp [2013], Wilson et al. [2014], Prabhu et al. [2011], Hannay et al. [2009]).

When one thinks of scientific software, one might first think of commercial or open source programs with thousands of users, such as Matlab, ImageJ, Imaris, etc. But there is a much larger set of programs that are created by small groups to perform domain-specific analyses. These small but critical programs are usually written in individual labs (McElreath [2023], Culina A [2020]), with a) narrow testing, b) no external review, and c) poor documentation. Labs spend weeks or months re-creating these small programs based on the literature, but they are hard to reuse because they often d) make assumptions about the format and organization of the data. As there are infinitely many ways to write a good computer program, there are barriers to e) understanding each new program and f) gaining trust that the program does what it claims to do. These problems are understandable, as the graduate students and postdocs who create these programs are focused on publishing results and getting their next job instead of writing a piece of software. While code is usually shared openly on sites like GitHub, these barriers have meant that, realistically, most programs are not reused or even read.

Despite good intentions and some internal efforts at testing, it is easy for calculations to go awry. In one case, a bug in custom software introduced errors into the structure of the ATP-binding cassette transporter MsbA that led to the retraction of 5 structural biology papers, including 3 in *Science* (Chang et al. [2006], *Miller [2006]). A similar error in in-house software found false evidence of a relationship between synapses and orientation columns in the developing visual cortex that was subsequently retracted in Neuron* (Katz et al. [1998]). A bug in common fMRI pipelines produced false-positive errors over a period of about 15 years (Eklund et al. [2012, 2016]). In a famous case, an economic analysis that in part caused some European countries to adopt austerity measures was found to have been in error because a spreadsheet calculation accidentally omitted data from some countries (McElreath [2023], Herndon [2014]). Use of standard spreadsheets as analysis tools has introduced widespread errors into gene names in genomic studies (Zimmerman [2014]). The consequences of limited testing must be worse than we understand because many errors are likely never uncovered.

The creation of common file formats like NWB (Rü bel et al. [2022]), BIDS (Gorgolewski et al. [2016, 2017]), OpenEphys (Siegle et al. [2023]), and data interfaces like Spyglass (Lee et al. [2024]) or NDI (García Murillo et al. [2022]) help with data management but do not solve these domain analysis problems by themselves. However, they do allow the creation of analysis applications that do not need to be concerned with the organization or underlying file format of the raw data. In this paper, we will take advantage of NDI, which creates a platform-independent, FAIR (Findable, Accessible, Interoperable, and Reusable) Wilkinson et al. [2016], searchable database of documents containing all raw data and analyzed data.

We have developed an applet motif that we call *calculators* that provides a framework for building robust, tested, and easily reusable analysis pipelines. *Calculators* greatly constrain the design and documentation choices for the developer, so that the code that performs the meat of the analysis and the documentations are in places where users and developers expect, greatly simplifying learning and adoption. Here we showcase the motif by developing *calculators* for vision science that perform the common actions of analyzing orientation and direction selectivity (Carandini and Ferster [2000], Ringach et al. [2002], Wu and Gazelle [2025]), spatial and temporal frequency selectivity (Movshon et al. [1978], Hawken et al. [1996], Heimel et al. [2005]), speed tuning (Priebe [2006]) and other forms of interdependence of spatial frequency and temporal frequency tuning, contrast tuning (Peirce [2007]), and Hartley stimulus reverse correlation (Ringach [1997]). These *calculators* have been used in published papers (Griswold and Gazelle [2025]) and preprints (Casanova et al. [2025]) and are being applied to our works-in-progress. We discuss the design principles of *calculators* and their advantages and disadvantages for reliable domain-driven computation in fast-moving scientific projects.

## 2 MATERIALS AND METHODS

The *calculator* motif idea can be implemented in any language, but we used MATLAB (MathWorks, Natick, MA) versions 2019b-2025a. The *calculator* base class was added to the Neuroscience Data Interface (https://github.com/VH-lab/NDI-matlab) and the visual neuroscience calculators are in a separate repository (https://github.com/VH-Lab/NDIcalc-vis-matlab). Users can write their own *calculator* objects in their own repositories by following the motif.

### 2.1 General methods for fits

For *calculators* that calculate response fit and index values, the raw responses were preprocessed so that the response to a control stimulus was subtracted, so that the responses reflect the activity of the neuron due to the stimulus (rather than any ongoing background activity).

The fit *calculators* compute two measures of statistical significance of the responses. First, they compute a *visual responsiveness p-value* by calculating an ANOVA over all the stimulus presentations and the control stimulus. This estimates the likelihood that there is no statistical variation across all of the stimuli and the control stimulus. Second, they also compute an *across stimulus anova p-value* by calculating an ANOVA over all the stimulus presentations (not including the control stimulus). This estimates the likelihood that there is no statistical variation in the responses across all the stimuli.

### 2.2 Orientation and direction tuning curves

The selectivity of each neuron was quantified using one minus the circular variance [Ringach et al., 2002, Mazurek et al., 2014]. For orientation selectivity:

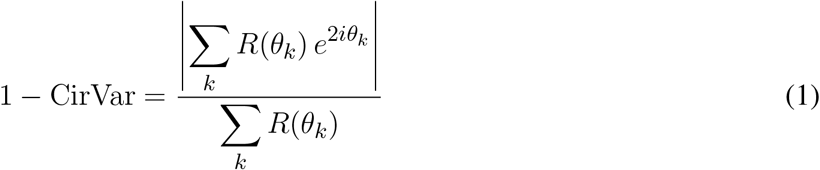

and for direction selectivity:

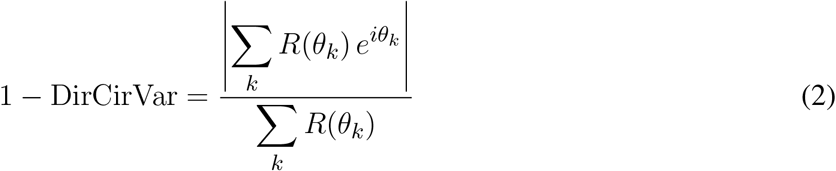

where *R*(*θ*_*k*_) is the mean response to a stimulus moving in direction *θ*_*k*_, and the sums are taken over all *k* stimulus directions. Both measures range from 0 (untuned) to 1 (perfectly tuned). The factor of 2 in the exponent of the orientation measure accounts for the 180° periodicity of orientation, such that responses to opposite directions contribute equally to the orientation selectivity estimate.

The data are also fit to a double Gaussian function of the form:

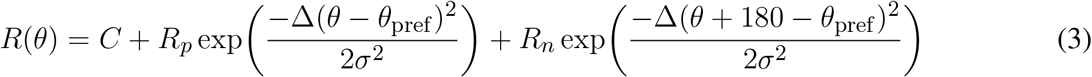

where *R*_*sp*_ is the untuned response of the neuron, *R*_*p*_ is the above-offset response to the preferred direction, *θ*_pref_ is the stimulus angle that evokes the maximum response, *σ* is the tuning width parameter, *R*_*n*_ is the above-offset response to the null direction, and *C* is the offset constant. The exponential function is defined as exp(*x*) = *e*^*x*^, and Δ(*x*) is the angular difference function defined as Δ(*x*) = min(*x, x* − 360, *x* + 360).

### 2.3 Contrast

The `ndi.vis.calc.contrast tuning` calculator object computes three fits of contrast tuning given a set of stimulus responses at a range of contrast values as described in Naka [1966], Albrecht and Hamilton [1982], and Peirce [2007]. The three fit functions are modified Naka-Rushton functions of the form

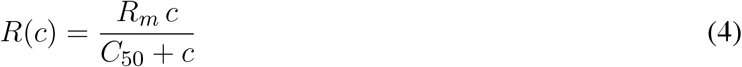

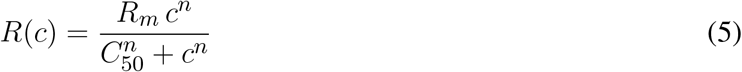

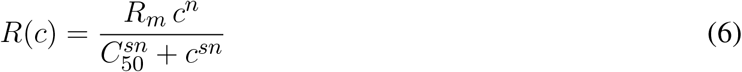

The calculator examines the raw data and the fits and makes determinations of the *empirical C50*. It also computes the *relative maximum gain*, the empirically biggest value of dR/dc for the fits or for the interpolated raw data, and, for the saturating equationPeirce [2007], the Saturation Index:

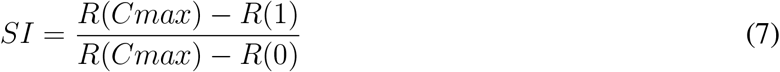

where *Cmax* is the contrast with the highest response *R*(*Cmax*).

### 2.4 Spatial and temporal frequency

There are two major types of indexes returned. One set are “fitless” index values that compute index values based on the empirical measurements without fitting. The second type are fit parameters and associated index values.

#### Fitless Measures

Several measures are computed directly from the empirical responses without fitting. The preferred frequency Pref is the frequency at which the peak empirical response occurred. The half-maximum points L50 and H50 are the frequency values below and above the peak, respectively, at which the empirical response dropped below half the maximum; if no such point occurred, L50 = −∞ and H50 = −∞. The bandwidth is defined as the interval between L50 and H50 in octaves,

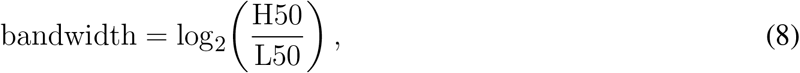

and is ∞ if either L50 or H50 is infinite. The low-pass index is the rectified response at the lowest tested spatial frequency divided by the rectified response at the peak spatial frequency, where rectification prevents responses from falling below zero; values range from 0 to 1, or NaN in the degenerate case of 0*/*0. The high-pass index is defined analogously using the response at the highest tested spatial frequency.

#### Difference of Gaussians

The data are fit to a Difference-of-Gaussians function of the form Heimel et al. [2005]:

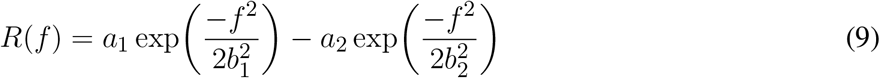

where *a*_1_ ≥ 0 is the amplitude of the first Gaussian, *b*_1_ is the fall-off of the first Gaussian, *a*_2_ ≥0 is the amplitude of the second Gaussian, and *b*_2_ is the fall-off of the second Gaussian.

#### Movshon Fit

The data are also fit to the function of Movshon et al. [2005], with or without a constant term *C*:

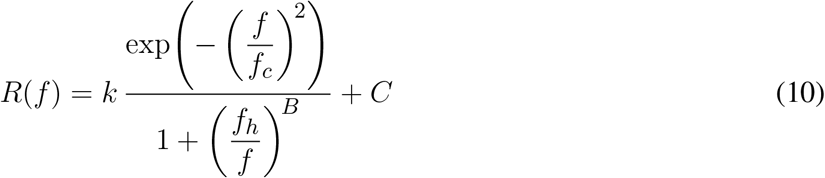

where *k* ≥ 0 is a scaling factor, *f*_*c*_ is the characteristic frequency, *f*_*h*_ is the corner frequency of the low-frequency limb, and *B* is the slope of the low-frequency limb. The constant term *C* is included in one version of the fit and omitted in the other.

### 2.5 Speed tuning

The data are fit to a two-dimensional speed tuning function as described in Priebe [2006]. The fit function is a modified two-dimensional Gaussian of the form

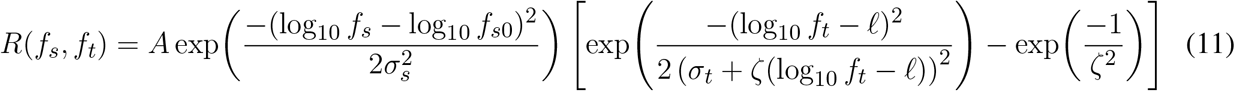

where

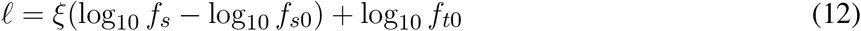

is the log temporal frequency at the preferred speed given spatial frequency *f*_*s*_. The fit parameters are determined by least squares. Here *A* is the peak response of the neuron, *ζ* is the skew of the temporal frequency tuning curve, *ξ* is the speed parameter relating preferred temporal frequency to spatial frequency, *σ*_*s*_ is the tuning width for spatial frequency, *σ*_*t*_ is the tuning width for temporal frequency, *f*_*s*0_ is the preferred spatial frequency averaged across temporal frequencies, and *f*_*t*0_ is the preferred temporal frequency averaged across spatial frequencies.

### 2.6 Hartley reverse correlation

The one *calculator* in this set that is not a fit function is the `Hartley` *calculator*, which computes the reverse correlation of responses of a neuron with a Hartley subspace stimulus (Ringach [1997]). The *calculator* returns a space-time-receptive field.

## 3 RESULTS

### 3.1 Data interfaces and databases allow a new kind of analysis pipeline

Many branches of science and neuroscience have domain-specific problems. A researcher studying gustatory cortex may want to examine stimulus selectivity to tastes; a researcher studying central pattern generators may want to understand how the period of an oscillatory is influenced by neuromodulators; and a researcher studying epigenetics may want to understand how accessibility of chromatin is altered by the chemical environment of an organism (**Fig. 1A**) . At their core, these domain-specific analyses comprise reading some primary data, performing a calculation, and storing the result so that it can be reported or be used for subsequent calculation.

**Figure 1.**
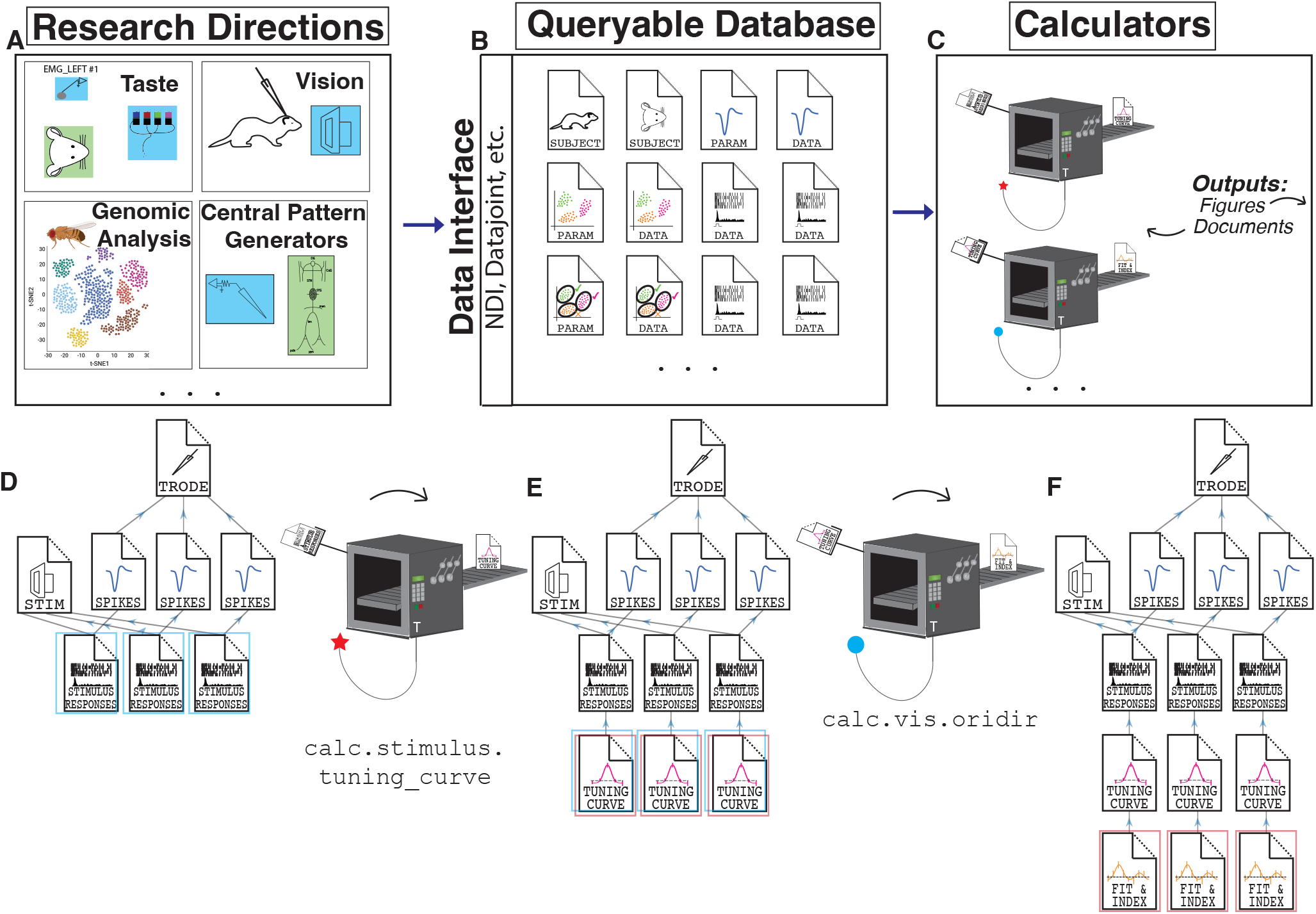
*Calculators* are a framework for creating domain-specific computations for a diverse array of disciplines that operate on and store their results in a database. **(A)**: Examples of specific domains in neuroscience: taste, vision, central pattern generators, genomics. Experiments from these domains produce data that can be stored in a **(B)** standardized, queryable database such as NDI or DataJoint. Such a database contains documents with information on the experimental subject, parameters, raw data, processed data, and more. **(C)** *Calculators*, which will be unpacked further in the next figure, perform analyses by searching the database for potential input documents **(D)** and then return output documents of a specific type. These documents are then added to the database **(E)**. By using a set of *calculators*, one creates *pipelines*. The next *calculator* searches documents to operate on **(E)**, performs its calculation, and adds the results back to the database **(F)**. This search-and-add system removes the need for explicit wiring of input documents to output documents. Instead, analysis pipelines emerge from multiple *Calculators* searching for successive document types on which to perform analyses.

Historically, all of these stages have offered barriers to good testing, readability, and reuse. The primary datasets themselves have been collected using a myriad of file formats and innumerable styles of organization. But now, modern approaches using inclusive file formats such as NWB (Rübel et al. [2022], Teeters et al. [2015]) or frameworks such as NDI (García Murillo et al. [2022]), DataJoint (Yatsenko et al. [2015]) allow analysis tools to query experimental data and analyses, agnostic of the experimental setup or underlying file systems. The availability of formats and systems that standardize data access now allows solutions that focus on the calculation and sharing side of the problem.

A key building block of our *calculator* motif is the use of typed databases. Interfaces like NDI and DataJoint and formats that offer rigorous extensions like NWB allow data elements to be expressed as a database (**Fig. 1B**). *Calculators* (**Fig. 1C**) search the database for data documents or sets of data documents upon which it can perform its calculation (**Fig. 1D**). In our example figure, a tuning curve builder searches the database for stimulus responses of a particular type. Then, it stores the result of each calculation as another typed data document, and adds each document to the database (**1E**). In our example, there is a document that stores the data for the tuning curve and it is added to the database. Therefore, the database of results documents grows. To perform a calculation that builds on top of another calculation, such as creating a fit to a tuning curve, another *calculator* that searches for tuning curve documents is run, and it finds documents that it can operate on. Then, the fits are calculated and placed in typed documents that are once again added to the database (**Fig. 1F**). The database of results keeps growing. Because the database is full of well-defined, typed documents, it can be shared and read by other researchers. In this manner, a set of *calculators* comprises an analysis *pipeline*.

### 3.2 The motif

The `calculator` motif is designed to make domain-specific computations easy to create, understand, and re-use. A graphical representation of the motif is shown in (**Fig. 2**). *Calculators* search for database documents that they can operate on (**Fig. 2A**) and perform a stereotyped computation (**Fig. 2B**). In order to showcase its behavior to potential new users and to gain their trust, the *calculator* must be capable of generating self-tests that it can display graphically (**Fig. 2C**). The documentation for the *calculator* is found in a consistent location (**Fig. 2D**) on disk and in a graphical user interface. A key component in the flexibility of *calculators* is their ability to take input parameters (**Fig. 2E**) to modify the calculation. This capability allows different instances of the same *calculator* object to perform different actions (for example, the tuning curve *calculator* might look for varying stimulus orientation in one instance, and look for varying stimulus tastants in another instance). To simplify the actions of a *calculator*, a *calculator* object can only produce a single type of output document whose form is rigorously specified (**Fig. 2F**). Finally, each *calculator* must have an ability to make a plot of this document type (**Fig. 2G**). The user might make other plots, but each *calculator* must have a standard method for visualization.

**Figure 2.**
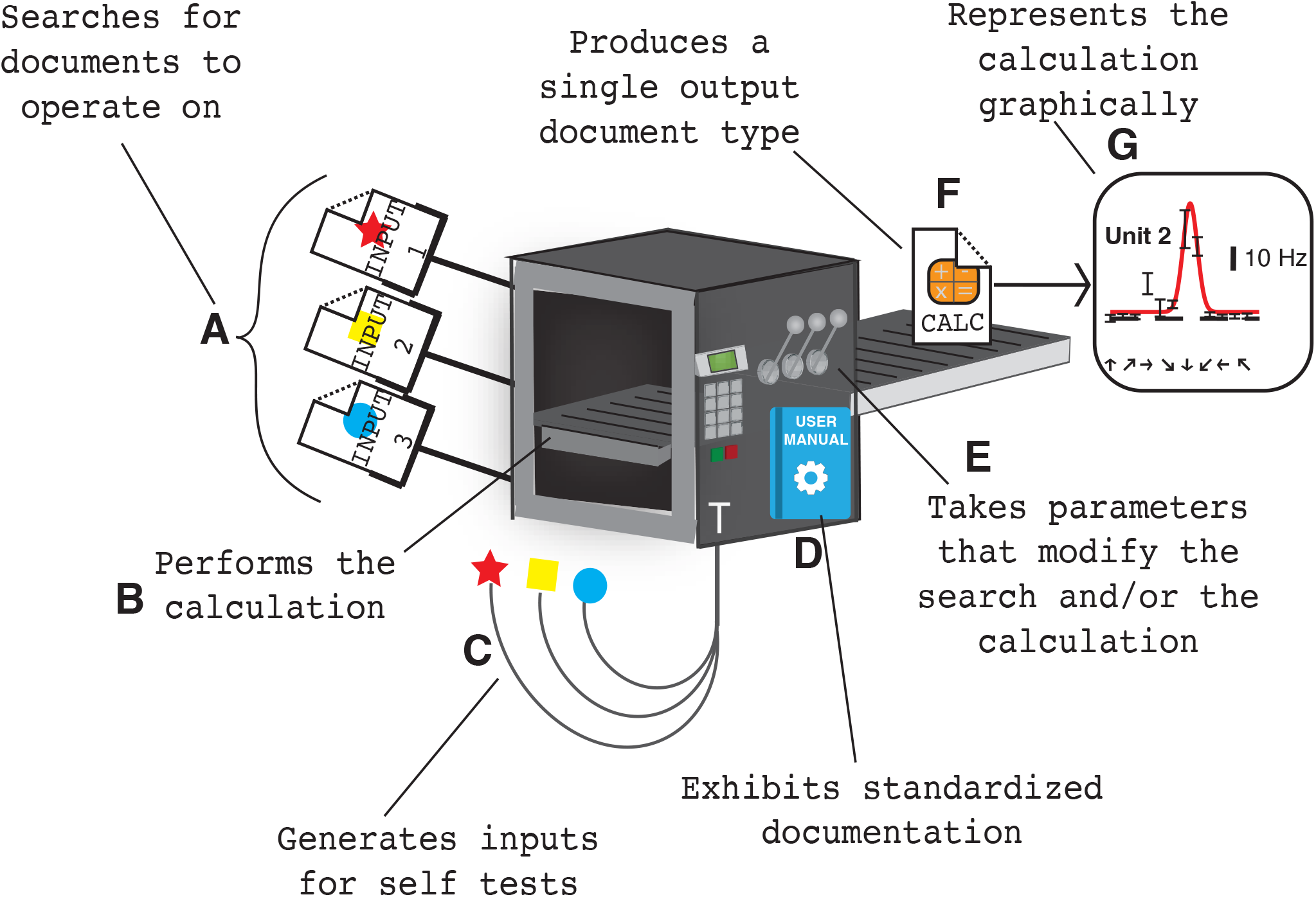
*Calculators* are designed to support broadly-scoped and reproducible analysis code. This is a graphical representation of the properties of all *calculators*. **A)** *Calculators* search through a database for default or user-specified document types, as indicated by the arms at the left seeking document sets of certain types. This makes it easier to reuse *calculators* in different scenarios since explicit wiring of inputs is not required each time. **B)** Once proper documents are located, they are used as input for analysis (into the box for computation). **C)** *Calculators* must be able to generate input documents for self-tests (symbolized by typed input arms that aren’t real documents). Testing on a sufficient variety of possible inputs provides trust in analysis results and an appreciation for the range of options the *calculator* can handle. **D)** *Calculators* provide standardized documentation to promote their re-usability (denoted by user manual on the box). **E)** *Calculators* take parameters (levers and buttons on the box) to provide users with a way to adjust how the input search is carried out or how the analysis is performed. This provides flexibility for different user requirements. **F)** Upon completion, *Calculators* each produce one document of a specific output document type (document coming out of the box), providing a standard that downstream analyses can count on. **G)** *Calculators* are required to provide a default plotting method (figure) to provide visualization of the output document to enhance trust in the results. The user might use this method or their own method for their own work but calculators must provide a default.

To build a *calculator* object, the developer follows a template and needs only to write a schematic of the output file format and a few methods (functions of the object) (**Fig. 3A**). One must write a document schema for the output document type, and specify this document type in the *calculator*’s constructor method (function). Furthermore, one must write a `calculate` method that actually performs the calculation on the input documents and produces the output document. One specifies the default search parameters that help the *calculator* find its inputs in two short methods (functions) called `default_search_for_input_parameters` and `default_parameters_query`. Finally, one must include code for plotting (`plot`) and documentation to describe a) the calculation, b) the output document type, and c) how to specify and adjust the search parameters to vary the *calculator*’s behavior.

**Figure 3.**
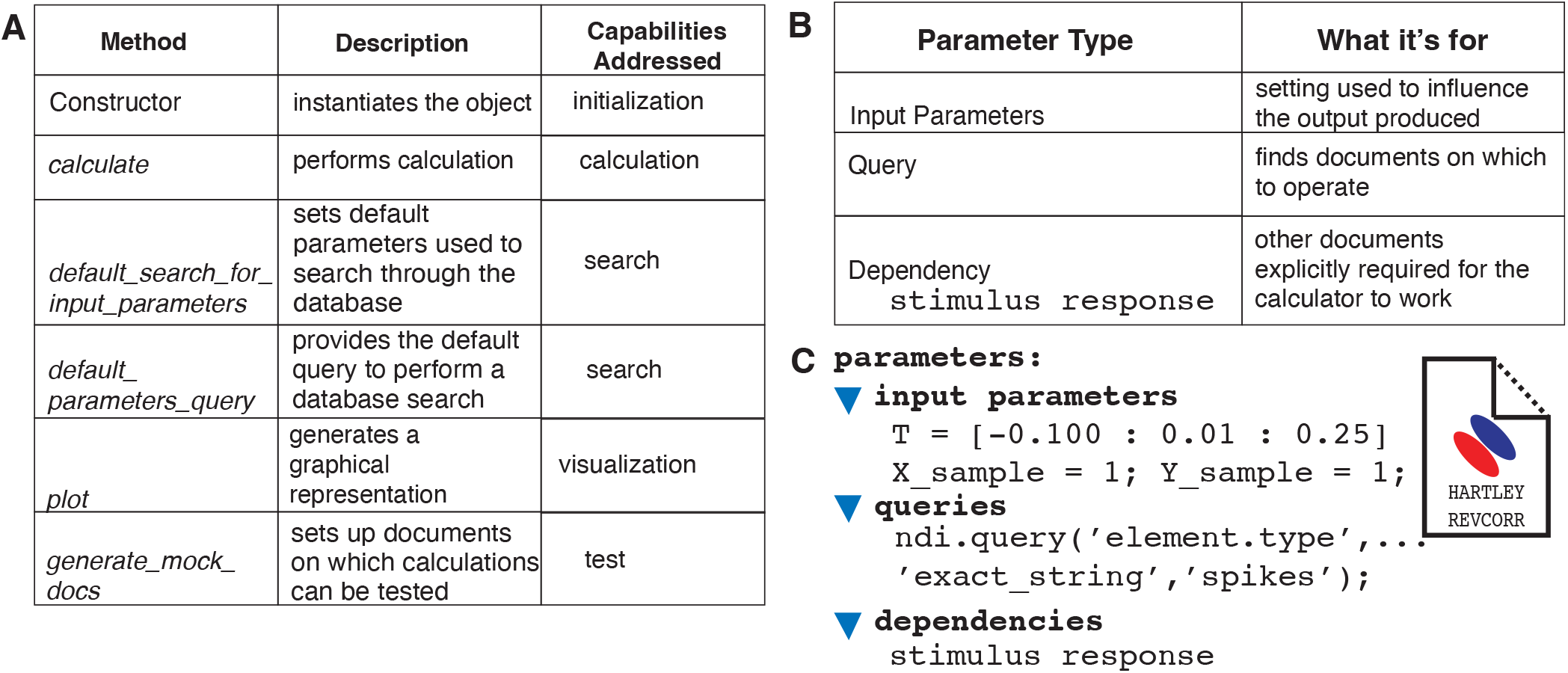
Software components of a *calculator*. **A)**. Required methods (function) of all `ndi.calculator` objects. Customizing these methods endows each class which its unique behavior. **B)** *Calculators* are driven by a parameter object that includes three components. Input Parameters are a set of names and values that alter how a *calculator* performs its computation. Queries describe how the *calculator* finds its input documents. Dependencies list specific document identifiers that the *calculator* should use to perform its calculations, and is usually populated by the query objects. **C)** Example of a set of parameters for the `ndi.calc.vis.hartley` object. The editable input parameters include the time resolution (stimulus time offsets) at which to calculate the reverse correlation and whether the answer should be binned in X or Y. The query restricts the elements to be operated on to spiking neuron elements.

A key source of the *calculator*’s flexibility comes from the ability to use different input parameters (**Fig. 3B and C**). In the input parameters, one can specify a variety of attributes about the documents to be found. To give an example, the input parameters of the `ndi.calc.tuningcurve` object allows wide variety of behaviors. One can specify the independent variable name(s) to dictate which stimulus parameter(s) should be used to identify stimuli to be included and the values of the stimulus parameter(s). The *calculator* can build N-dimensional tuning curves and include only stimuli that are a part of a particular set where certain parameters vary or have a particular value. For the Hartley reverse correlation reconstruction, one specifies the extent and resolution of the reconstruction in time in the parameter value T and allows the user to spatially bin in X and/or Y (**Fig. 3C**). Other *calculators*, like the `ndi.calc.vis.oridir tuning` have no adjustable input parameters and the calculations are made the same way each time.

*Calculators* produce a single, typed output document like the example shown in **Fig. 4**. Fields such as `app` hold information about the version of the program that generated the document, and `depends on` indicates the documents that were used in the calculation of the index and fit values. Finally, several fields of domain-specific computations about response significance and, for orientation tuning documents, vector- and fit-based analyses are reported. The document is stored in the experiment database and all fields are exposed to search.

**Figure 4.**
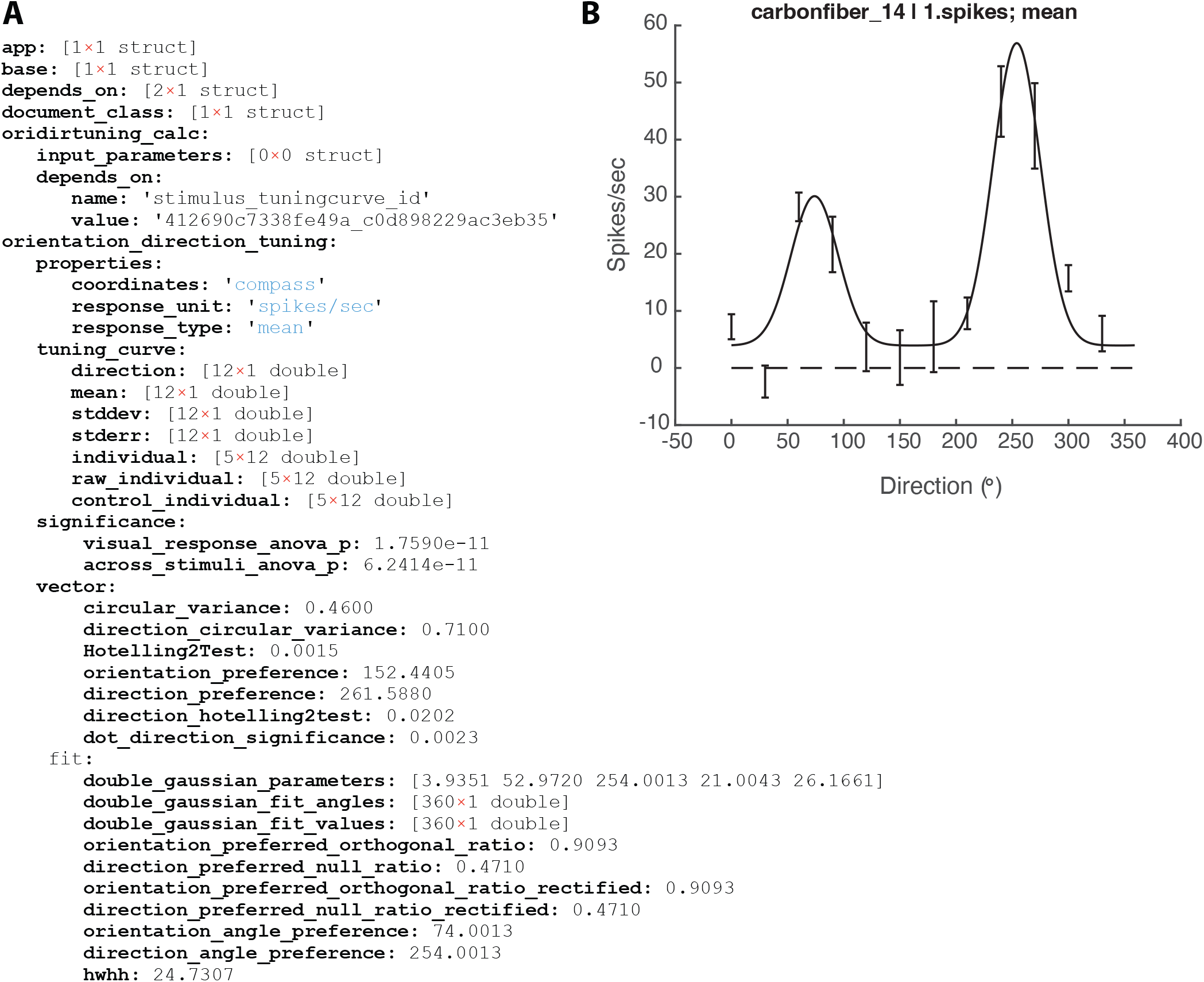
Example document output of a *calculator*. **A)** Fields list for documents generated by `ndi.calc.vis.oridir tuning`. The domain-specific fields are expanded showing properties, tuning curve parameters, significance evaluations of the responses, and parameter and index values derived from both vector analysis methods and fit-based methods. **B)** The default figure produced by the calculator for the same cell.

The *calculator* framework therefore allows complex pipelines to be specified with only a few elements. A graphical illustration of such pipelines are shown in **Fig. 5**.

**Figure 5.**
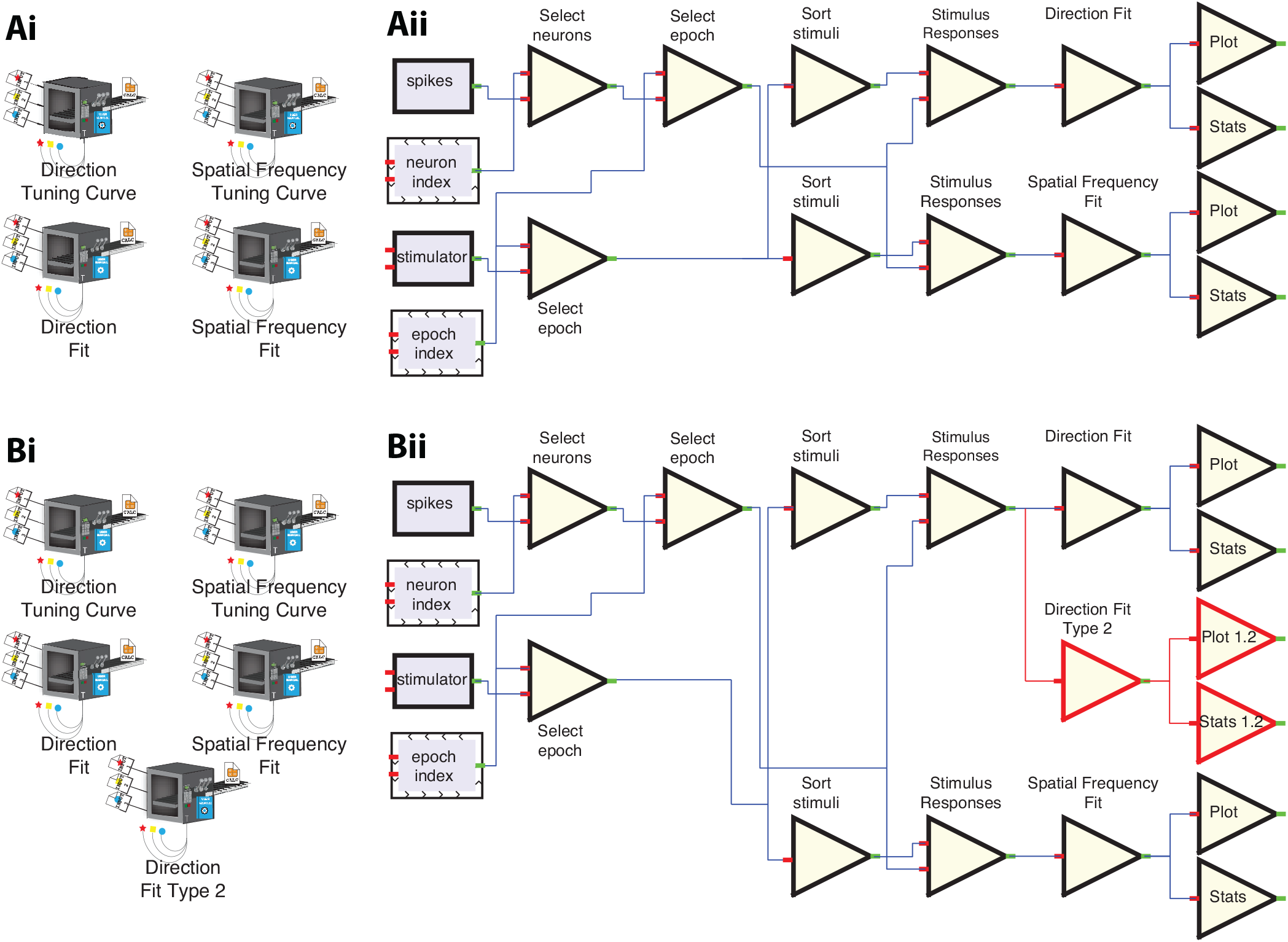
Visual illustration of NDI *calculator* pipelines. **Ai)** A set of NDI *Calculator* instances for creating tuning curves and for creating fits to tuning curves, in this case, for stimulus Direction and stimulus Spatial Frequency. The *calculators* that create tuning curves search the database for stimulus responses that match their parameters and create tuning curves. Then, the fitting *calculators* search for tuning curves that match their parameters and create fits and plots. **Aii)** Wiring diagram equivalent. Basic NDI functions produce stimulus responses for all stimulus presentations and all spiking neurons. The system loops over all these combinations of stimulus presentations and spiking neurons (represented by the arrowhead border in the neuron and epoch index boxes) and computes stimulus responses. Then, stimulus responses are organized into tuning curves, which are then fed into fitting modules (Direction Fit, Spatial Frequency Fit), from which plots and fit statistics are produced. **Bi)** Adding a new calculator instance to a set changes the pipeline. Here, the user has added a second fitting algorithm that searches for tuning curves that vary in Direction. **Bii)** This new calculator instance, by virtue of searching for documents that match, pipes the tuning curves that vary in Direction through the new fitting algorithm, producing plots and fit statistics. The provenance of documents (flow) can be examined by examining the database (see Figure 1).

In all, a set of *calculator* instances allows complex pipelines to be specified. To help understand the structure of pipelines that are created by *calculators*, we illustrate the equivalent pipeline “wiring diagram” for two sets of *calculators* in **Fig. 5**. A pair of *calculator* instances that are set up to create tuning curves for different stimulus parameters is shown in **Fig. 5Ai** along with two other *calculator* instances that compute fits of direction or spatial frequency tuning curves. In code, there would be 3 object types written, but by using different parameters for the tuning curve *calculators*, one creates two instances (and writes no code to create the second instance). In the first layer (**Fig. 5**), NDI loops over all combination of stimulus presentations, neurons, and their respective recording epochs to produce stimulus responses. The tuning curve *calculators* search over this space and find all combinations that they can sort into tuning curves (“Sort Stimuli” triangle). Then, these tuning curve response sets are found by the Direction Fit and Spatial Frequency fit *calculators*, which produce documents that contain statistics and allow plotting.

Adding a new branch to the pipeline can be as simple as adding a new *calculator*. In **Fig. 5Bi**, a second Direction Fit type has been added. Because this calculator type searches for direction tuning curves, the extra calculations are automatically performed (**Fig. 5Bii**), enriching the database.

### 3.3 Example applications

We developed several *calculators* for our lab’s own workflows in vision science. All the fitting *calculators* provide some measure of statistical significance of the responses by computing an ANOVA across all stimulus conditions and the blank (answers if the cell is visual responsive) and a second ANOVA taken only across all stimulus conditions (answer if there is any significant variation across the stimulus responses).

A variety of vector-based (that is, fitless) (Ringach et al. [2002], Mazurek et al. [2014]) and fit-based approaches (Swindale [1998], Carandini and Ferster [2000], Mazurek et al. [2014] have been developed to quantify the orientation and direction selectivity of neurons in the visual system, although the same measures could be applied to other directional neurons such as head-direction cells. The `ndi.calc.vis.oridir` *calculator* returns 19 measures (**Fig. 6A**), including the circular variance in orientation or direction space, the preferred orientation or direction using vector methods, the coefficients of a double-Gaussian fit, and a variety of orientation and direction selectivity index values. This *calculator* has been used for 3 papers from our lab (Reikersdorfer et al. [2021], Griswold and Gazelle [2025], Casanova et al. [2025]). Further, this *calculator* is being used in a collaboration on another lab’s data, with different formats and organization, that is being read with NDI, showing that *calculators* can readily analyze cross-lab data.

**Figure 6.**
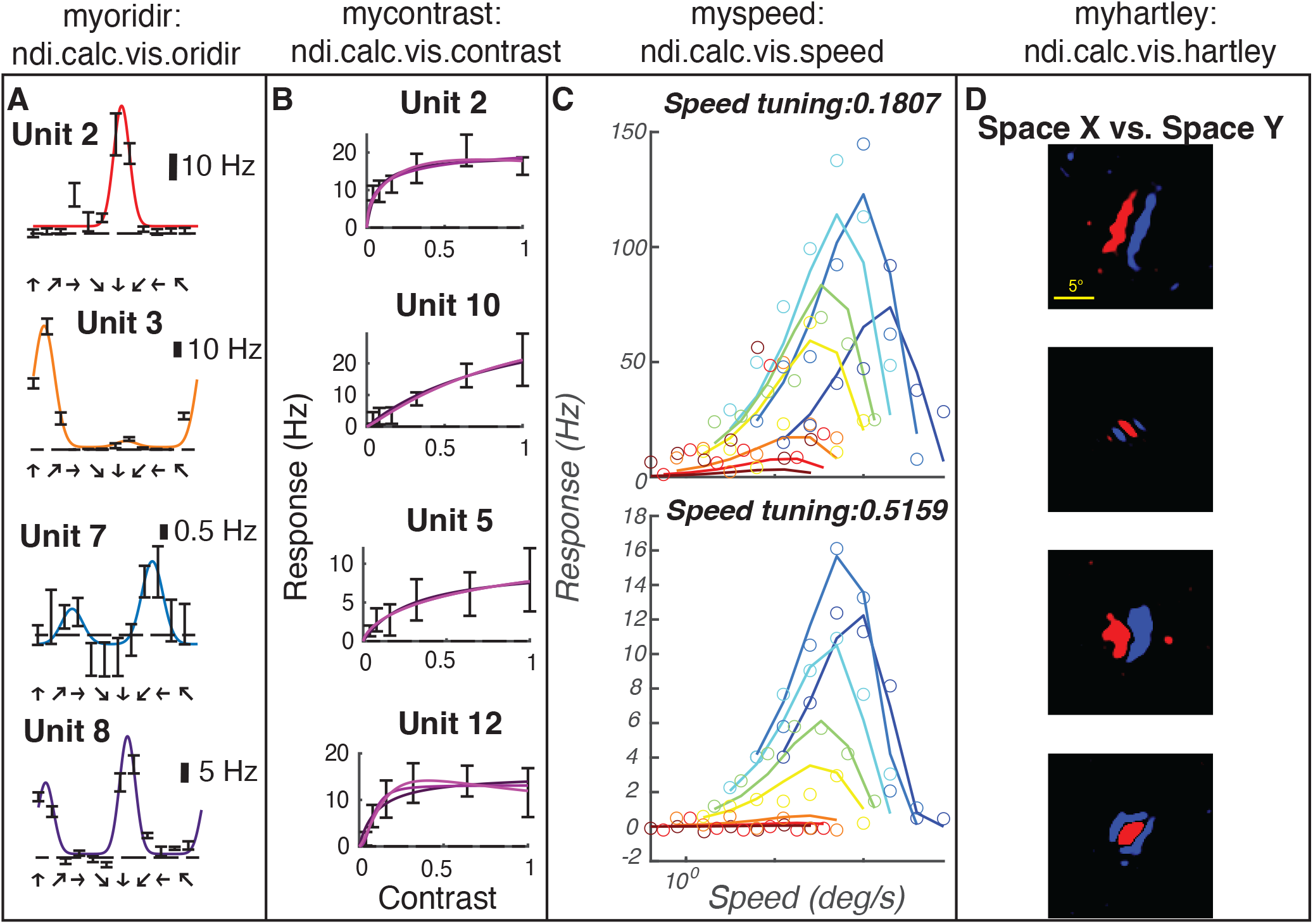
Examples of NDI *calculator* objects that form a visual analysis processing pipeline. **A)** ‘myoridir’ calculator instance of `ndi.calc.vis.oridir` generates analyses of tuning curves for direction based on vector methods and a double-Gaussian fit. **B)** ‘mycontrast’ *calculator* instance (type `ndi.calc.vis.contrast`) generates fits and index values for contrast tuning curves based on Naka-Rushton fits (conventional, exponential, and saturating forms). **C)** ‘myspeed’ *calculator* instance generates fits to 2-dimensional tuning curves of responses to stimuli that co-vary in spatial and temporal frequency, allowing calculation of speed tuning index values. **D)** ‘myhartley’ *calculator* instance generates receptive fields from reverse correlations with Hartley stimuli. RED indicates ON responses while BLUE indicates OFF responses.

In the domain of stimulus contrast, most researchers apply equations used by Naka and Rushton (Naka [1966]) with various extensions for increased slope or saturation (Peirce [2007]). The `ndi.calc.vis.contrast` *calculator* returns three Naka-Rushton-style fits (**Fig. 6B**) as well as empirical measurements of the 50% response point, the peak response, the relative maximum gain, and saturation index. This *calculator* has been used in one published paper (Griswold and Gazelle [2025]).

For spatial and temporal frequency, we compute several fit measures including a standard difference-of-Gaussians model as well as a bandpass model due to Movshon and colleagues (Movshon et al. [2005]). For the empirical data and for the fits, we calculate the frequency with the peak response, and search lower and higher frequencies to find the (interpolated) frequency where the response drops to 50% of this maximum value (low frequency cut-off and high frequency cut-off). In all, N quantities are calculated. These *calculators* have been used in one published paper (Griswold and Gazelle [2025]).

We also have a *calculator* called `ndi.calc.vis.speed` (**Fig. 6C**) that computes responses to joint measurements of spatial and temporal frequency responses and is capable of uncovering relationships where temporal frequency tuning depends on the spatial frequency of the stimulus or vice-versa (Priebe [2006]). This fit helps determine if neurons are tuned to speed *per se* or rather if the cell is just tuned a fixed spatial frequency that does not vary with the temporal frequency of the stimulus (independent tuning). This *calculator* has been used in one preprint (Casanova et al. [2025]).

Finally, we have a calculator for computing reverse correlation of neural responses with Hartley subspace stimulation (Ringach [1997]) which we have used in unpublished work to measure space-time receptive fields of visual cortical neurons in the ferret (**Fig. 6D**).

### 3.4 Graphical user interface

To translate the conceptual framework of *calculators* utilizing NDI into a practical tool for daily research, we generated a corresponding MATLAB Graphical User Interface (GUI). Here, we provide a user interface program built to serve as a bridge between the NDI database and the researcher to make robust and standardized analysis accessible to naive programmers. This interface is designed to directly expose the core benefits of the *calculator* architecture: modularity, reusability, and independence from pre-wired pipelines. The GUI provides a template for *calculator* parameters so that users can easily create *calculators* and pipelines for their domain of research and intended analysis. Instead of requiring users to write code to connect analysis steps, the GUI provides an integrated environment for dynamically searching the database for input documents, configuring parameters for one or more *calculators*, and executing entire analysis sequences as a pipeline. This approach effectively streamlines the process of building, testing, and running complex analyses, making the system accessible to users with varying levels of programming expertise.

The GUI provides a single interactive tool to consolidate the functions of NDI *calculators*. Users are provided with documentation of a *calculator* function, alongside default parameter code, that can be easily edited based on the provided template and tutorial. This allows for user-friendly customization of NDI *calculators* and input parameters, which builds reliable and reusable analysis pipelines. This tool can be utilized for both rapid exploratory data analysis and reproducible production of figure-ready data. For detailed guidance on implementing the GUI, refer to the provided NDI *calculator* GUI User Manual. An example user workflow is presented here.

A user loads the GUI Pipeline window in MATLAB (**Fig. 7A**). Here, they can either select an existing pipeline or create a new pipeline. This pipeline consists of *calculators* that are run in succession, although the order is unimportant due to the fact that *calculators* search the database for their input documents. *Calculators* can be added or removed from the pipeline based on the desired output of the analysis pipeline. For example, if one wishes to analyze data to produce orientation tuning curves, they will include the `ndi.calc.vis.oridir` *calculator*, which they can nickname whatever they want for user customization. Pipelines can be customized with different *calculators* so that the GUI can flexibly handle different types of data and analyses. Within the pipeline window, the user can also select an existing NDI data session (S,**Fig. 7A**) to which they wish to apply the pipeline. Outputs of the *calculators* within this pipeline will be saved as NDI documents, or produce figures.

**Figure 7.**
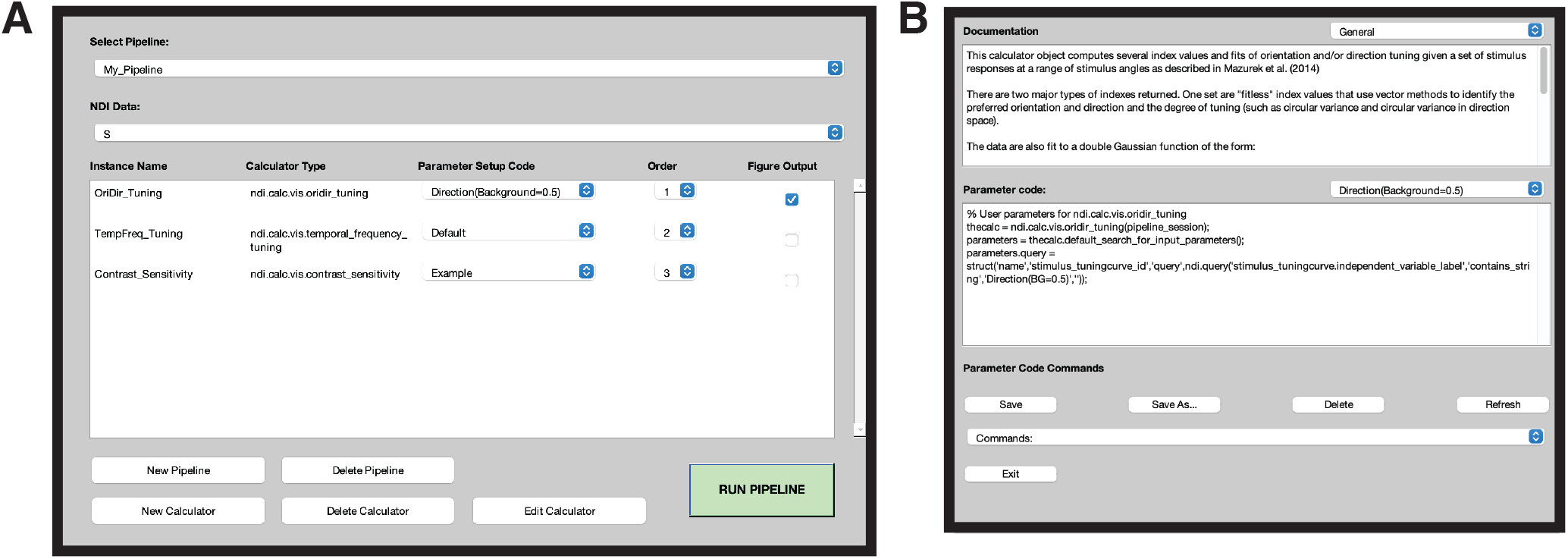
MATLAB graphical user interface to run, edit, and create pipelines of NDI *calculators* for neuroscience data analyses. **A**: Main GUI window for creating customized pipelines of *calculators* for a desired analysis. **B**: NDI *calculator* editing window to edit parameters for customized implementation and view *calculator* documentation.

The GUI Pipeline window also allows the user access to edit the included *calculators* (**Fig. 7B**). When editing, the user has the option to further edit the parameters of a *calculator*, changing the output of the analysis based on the particular needs of the experimenter. In this Editing window, the user is presented with the *calculator* documentation, and the parameter code which is modifiable. If the user alters the *calculator* parameters, they can save this as a new parameter template, available for later utilization or sharing. They are also able to run various commands, such as searching for input or output documents, to explore the *calculator*.

Overall, with this GUI the user is able to flexibly implement many different analysis pipelines which quickly and reliably produce figures for a variety of neuroscience experiments, with very little programming experience. The provided default parameter template provides a framework for a naive user to alter *calculator* parameters, so they can build desired analyses through the GUI. Pipelines can be easily built and executed to analyze a large amount of data in a variety of ways through interaction with a single interface.

### 3.5 AI Agent manual

AI coding agents have emerged since we have begun this project, and we have found that AI agents such as Gemini and Claude do a good job at writing a draft for new *calculators* while conforming to the structure of the motif. We provide a manual for agents in the GitHub repo that has been used successfully to create calculators for ongoing work in the lab. All of the *calculators* reported in this work in the domain of vision science were created by hand.

## 4 DISCUSSION

We have described a motif for performing domain-specific analyses that brings some common practices of software development to bear in order to improve rigor, robustness, and reuseability. The documentation and key computing code are in consistent places, allowing new users to gain familiarity quickly. *Calculators* must demonstrate end-to-end test capability to help users build trust that the *calculator* does what it claims to do. *Calculators* can be composed into flexible pipelines that exhibit standardized and searchable output at each stage, helping scientists working even in small domains conform to FAIR principlesWilkinson et al. [2016].

### 4.1 Domain specificity

Scientific research consists of some broad classes of data that enable the development of robust and well-tested pipelines, such as the domains of spike sorting for electrophysiology data (e.g., KilosortPachitariu et al. [2024], SpikeInterfacesBuccino et al. [2020], the Plexon Sorter, JRClustJun et al. [2017]) or cell finding for calcium imaging (e.g., Suite2pStringer et al. [2026], EXTRACTDinç et al. [2024], CaImAnGiovannucci et al. [2019], or NANSEN).

Further, most scientific works depend on the development of small pieces of software that may only be applicable in specific domainsHerndon [2014], McElreath [2023], Huang and Lapp [2013], Wilson et al. [2014], Prabhu et al. [2011], Hannay et al. [2009]). While one can post the code on repositories such as GitHub, there are substantial barriers for others to try to understand or reuse the code because of the infinite flexibility of coding languages. While there are many general purpose pipeline applications, such as Snakemake, Nextflow, and KNIME, these tools primarily solve the dependency problem (making sure the applications run in order) rather than addressing the software engineering culture issue of including documentation in certain places, performing testing, and standardizing outputs.

The *calculator* motif narrows the choices that the developer can make in the interest of consistent readable, documentation, testing, and output data structures, to guide scientific programmers into using good practices and to increase the odds that the code could be picked up and used by other groups.

### 4.2 FAIR development

The *calculator* motif produces standardized, searchable document types that contain the provenance of the documents that were used to build the computation, so the output satisfies all of the elements of FAIR: Findable, Accessible, Interoperable, and Reproducible Wilkinson et al. [2016]. The output can be searched with the appropriate query language (such as `ndi.query` for NDI) on personal computers or in the cloud as sites like NDI Cloud. Further, pipelines build with *calculators* are FAIR at every stage; each intermediate step in the pipeline produces a standardized product.

### 4.3 Scalability

One nice feature of using data interfaces with databases is that the analyses can be scaled arbitrarily. One can run *calculator* pipelines on very large databases and spread the computation among as many parallel systems as one wishes, and add the results back to the database. Once you have a *calculator* and a pipeline, it takes the same effort to ask the system to analyze one item or millions of items.

### 4.4 Time burden

Once accustomed to the process, making a calculator does not take too long, and AI coding agents are good at writing a decent first draft. However, it can take much longer than the “hacking”-style analysisMcElreath [2023] that the senior author has done for most of his career. Because the functions will have a longer life, one spends more time revising and refining a calculator than a standard analysis function. Often, if one has not employed a particular analysis previously, there is a learning curve as the function develops the sophistication and resilience necessary to be useful in the analysis and avoid the pitfalls (such as local minima) for the project at hand. But it takes more effort to then consider how the calculation should work in the general case, and to anticipate and solve potential problems that future users may have that were not exposed in one’s specific project. It is clearly more rigorous and careful to spend this time, but whether this time would be better spent making new studies or new analyses is a challenging question. While there are no guarantees, the *calculator* motif provides a better chance that this effort will live on.

The structure of the motif also helps to ensure that AI coding agents write code that is well documented, well tested, and reusable. Because the structure introduces constraints and the output document introduces standards, the code that is generated by agents is easier to review to ensure that best practices have been followed and that the AI generates a component that interoperates with other software.

### 4.5 Testing: Problems solved vs. unsolved

The tests provided by *calculators* provide assurance against some but not all pitfalls in numerical analysis. The tests indicate that the computations do not suffer from machine precision errors or get stuck in local minima for the particular test cases shown. But these tests do not obviate the need to be careful in interpreting the data. For example, when responses are weak, many fit parameters become arbitrary, such as orientation preference angle or *c*_50_ of a contrast fit, and one still has to be cautious making interpretations or performing direct statistics on such parametersMazurek et al. [2014].

### 4.6 Wider adoption and reuse

If the concept of *calculators* gains traction in other groups over time, it is possible to establish peer-review communities of developers that can help test, validate, and improve *calculators*. Further, *calculators* can be cited in the methods sections of papers to credit the authors, and it is easy to generate RRIDs or other identifiers to make citing individual calculators easy.

The broader promise of the *calculator* motif is that it does not require the resources or community infrastructure that large shared software projects depend on to succeed. Such projects typically rely on institutional backing, grant funding, and large volunteer communities — conditions that are unlikely to exist for the long tail of domain-specific analyses that most scientists actually need. The *calculator* motif instead attempts to make good software practices the path of least resistance for a single developer working alone. By constraining the choices a developer can make, the motif aims to produce code that is documented, tested, and readable almost as a byproduct of following the template — lowering the bar for a future user or collaborator to understand, trust, and reuse the work without requiring a formal review process to certify it. If *calculators* accumulate over time into a shared library of domain analyses, formal mechanisms for credit and peer review — such as Journal of Open Source Software publications or registration with a Research Resource Identifier — could be layered on top. But the motif is designed to deliver value even before that community exists, by making it easier for one lab’s careful work to become another lab’s starting point.

